# Quantifying the size and healing of volumetric muscle wounds using 3D Slicer on CT scans in a canine model

**DOI:** 10.1101/2021.07.13.452185

**Authors:** Dale Larie, Gary An, Scott Johnson, Kelly Hoffman, Yoram Vodovotz, Stephen Badylak, Chase Cockrell

**Affiliations:** Department of Surgery, University of Vermont Larner College of Medicine; McGowan Institute of Regenerative Medicine, University of Pittsburgh; Division of Laboratory Animal Resources, University of Pittsburgh; Department of Surgery, University of Pittsburgh School of Medicine

**Keywords:** Volumetric muscle wounds, volumetric muscle loss, wound healing, Computed Tomography, 3-dimensional Computed Tomography, Wound Volume Quantification

## Abstract

Volumetric soft tissue and muscle wounds can arise from trauma or necrotizing soft tissue infection. Quantifying the size of these wounds can be challenging, as they often have irregular borders and contours and invariably involve skin loss. 3-dimensional Computed Tomography (3dCT) has been used to characterize the volume of numerous tissue structures, but these use cases invariably involve structures for which clear anatomic borders exist. This is not the case for volumetric soft tissue or muscle wounds, where the volume of the wound being assessed, which is actually a void representing the absence of tissue, does not contain an explicit border at the superficial surface. We present a method that allows quantification of the void size of volumetric muscle wounds using CT scans processed with the software package 3D Slicer. This quantification allows us to chart the progression of healing in such wounds with sequential scans. The development of a means to quantify wound size and healing rate is a necessary capability in order to assess the efficacy of potential therapeutic interventions aimed at enhancing healing of such wounds.

## Introduction

Necrotizing soft tissue infections and massive trauma can result in considerable soft tissue loss. These injuries involve skin, subcutaneous fat, fascia and muscle and can present considerable challenges to clinical care. In addition to treating the underlying cause of the tissue loss (i.e., severe infection or trauma) the management of the wound itself can often be painful and labor intensive. Such wounds produce considerable physiologic and metabolic stress and can be vulnerable to secondary infections. Current care strategies include repeated debridement to remove marginal/infected tissue, the use of negative pressure wound devices, and a range of topical agents in conjunction with different types of dressing materials [1, 2]. Development of new therapeutic agents or modalities should be guided by some quantitative metric that can determine if the proposed therapy actually increases healing rate. However, given their often irregular shape, the fact that there is a depth to the wounds, and that the wound represents an absence of tissue (e.g., a negative space or void), it is nearly impossible to clinically measure the actual amount of tissue loss in volumetric wounds. 3-dimensional Computed Tomography (3dCT) has been used to quantify numerous types of tissue structures, such as tumors or organ volumes [3, 4]. However, the fact that the void of a volumetric wound is unbounded along its superficial surface prevents the use of standard methods of calculating 3dCT volumes. Complicating this further in terms of collecting and quantifying sequential measurements, a necessary process in assessing the rate of healing of the wound, is the deformability of the soft tissue and variability in the orientation of the individual in the CT scanner. To date we are not aware of any method that quantifies the volume of a volumetric soft tissue-muscle wound that includes a skin deficit.

## Methods

### Data Generation

The data used for this project were multiple CT scans performed on a canine model of volumetric muscle loss (VML) injury [5]. The experimental protocol was approved by the University of Pittsburgh Institutional Animal Care and Use Committee. In brief, dogs were anesthetized. After aseptic preparation of the surgical site the skin was incised to expose the biceps femoris muscle. Approximately 75% of the muscle, including vasculature and nerve, was excised surgically, creating an approximately 10 cm long x 4 cm wide x 2 cm deep VML defect. Hemorrhage was controlled with cautery as needed. Dry Tefla non-adherent gauze dressings were applied and the animals allowed to recover. At various intervals spanning 1, 2, 4, 7, 10 and 14 days after wounding, the animals were re-anesthetized and repeat CT scans were performed.

### Data Extraction

The first step in attempting to analyze the wounds is extracting data in a useable format from the raw CT data. 3D Slicer is a free, open source, multi-platform software package that can be used to compile and render CT scan slices into a 3-dimensional model of the scan [6]. Given variation in positioning in the CT scanner at each imaging episode, the scans each have an arbitrary position and rotation in space, which was not consistent between scans of different days of the same dog nor consistent across scans of different dogs. In order to standardize the data extracted for analysis to provide a consistent coordinate system all scans of the same dog were aligned in space with the same position and rotation. This was done by a rotation/translation transformation generated and applied to each CT scan so that all scans of the same dog have the same position and rotation as an arbitrarily picked “master scan” of that dog. This can be done with 3D Slicer’s Image Guided Therapy (IGT) package, which generates transformation matrices using manually picked landmark points on each scan whose relative positions remain fixed across scans of the same dog. The “master scan” for a particular dog was chosen and the rendered image rotated such that the major and minor axes of the wound in the master scan form a plane parallel to the XY plane of 3DSlicer’s global coordinate system. Next, it was necessary to identify fixed landmarks for consistent orientation across individual scans. The femur was chosen as a fixed, rigid and consistent object for orientation. 3D Slicer has the ability to generate segmentations based on tissue density, allowing 3-dimensional renderings of only the bones or soft tissue. Using femur segmentation, points were picked manually and a transformation from the current scan to the master scan is generated. While the bones might be well aligned, the soft tissue of the dog’s leg is deformable which can mean that the surface of the skin and the precise rotation of the wounds are not consistent and might require manual fine tuning. After all scans of the same dog have been aligned, a single set of crop boundaries are picked so that they contain the entire wound with a buffer of healthy skin for all days of the same dog. Using this boundary set, all scans were cropped and data extracted from 3DSlicer using the built in python API as a 3-dimensional matrix where the data at each point (X,Y,Z) represents the density value measured by the CT scanner at that point.

### Data Analysis

There are 3 major analysis tasks to be performed. The first task is to detect the edges of the wound. The second task is to use the detected edges of the wound to determine the volume of the wound in order to calculate growth rate of tissue over time. The final task is to attempt to differentiate between scar tissue and muscle tissue based on the CT scan density readings.

#### Task 1 – Wound Edge Detection

Detecting the edges of a wound is a trivial task by eye but attempting to label the edges entirely manually would be inconsistent and add unnecessary human error. At the same time, detecting edges using a strictly programmatic approach is difficult and does not generalize easily to images besides the test image. The proposed method for this task is a hybrid method that eliminates human bias as much as possible but still leverages the simplicity of manual labeling. There are three major steps for this edge detection method, and they can be broken down into image processing, manual outlining, and automated edge detection.

The first step in image processing is taking the 3 dimensional matrix that represents the 3-dimensional rendering of the CT scan and converting it into a 2-dimensional map of the height (Z) at the surface of the wound at each point (X,Y). This 2-dimensional heightmap will be used for the remainder of the edge detection task and will be used for the tissue growth rate calculation task. Taking the divergence of the heightmap will highlight steep changes in height, a feature which is present at the edges of the wound. To filter out noise, the divergence map is convolved with a gaussian filter and values below a set threshold are changed to be 0. The gaussian filtered map is then convolved with a “sharpen” filter, which highlights boundaries of objects in images. The values in the twice filtered map are then converted from continuous values into binary values using another set threshold. This binary bitmap is the final data returned from the image processing step, and it highlights the boundaries of the wound as well as any other bumps or sudden changes in height on the heightmap.

The next step is manually outlining the wound. Using a 3-dimensional rendering of the wound as a reference, points that are outside the wound but before any outside noise are manually marked on the binary bitmap returned from the image processing steps, and a polygon is generated using the selected points as vertices. That polygon is used as a mask, and all points on the bitmap that do not fall inside the polygon are set equal to 0. As a result, the only data that remains should be a bitmap representing the contours of the wound.

The last step in wound edge detection is using the masked bitmap to outline the wound. Any polygon drawn on a 2-dimensional grid will have pairs of points with coordinates where one coordinate position is the same and the other is different. As a result, it is possible to sweep across that polygon and find pairs of points along the perimeter of the polygon. Using the masked bitmap, for each X coordinate the largest and smallest Y coordinates that have non-zero value represent a pair of points marking the top and bottom edges of the wound at that point. Sweeping across all X values in the image will generate a set of pairs of points that define the detected top and bottom edges across the length of the wound. Repeating this process but sweeping across the Y values will generate a set of pairs of points that represent the left and right edges of the wound. Combining the top and bottom edge pairs with the left and right edge pairs will fully define the detected edge around the circumference of the wound. Using these detected edge pairs, it is possible to draw a convex polygon that outlines the perimeter of the wound.

#### Task 2 – Wound Volume Determination and Tissue Growth Rate Calculation

Two methods were tested to try to calculate the volume of the wound such that the rate of tissue growth in the wound could be calculated. The key challenge is to approximate the superficial surface of the tissue void, thereby fully bounding the volume of the wound. The first method (Method 1) generates a flat plane across the edge of the wound and calculates the volume of the void from the base of the wound up to the plane. The second method (Method 2) interpolates a rounded surface using the healthy skin edges that represents how a “healthy” leg would look. These methods do not explicitly measure the growth rate of tissue, but instead implicitly measure the growth rate of tissue by measuring the change in the volume of the void left by the wound.

The first method projects a flat plane across the edges of the wound as the simplest surface that can be drawn to enclose the wound void. A plane can be generated by selecting points along the edges of the wound where the skin ends. Choosing an arbitrary number of points along the perimeter of the wound will generate a plane that approximates the average height and angle of the perimeter of the wound in 3-dimensional space. Summing the difference between the Z value of this plane and the height (Z) from the wound heightmap at each point (X,Y) within the bounds of the wound would produce the volume of the void.

However, there are intrinsic problems with this method. First, the surface of a dog’s leg is naturally curved. Using a flat plane to try to approximate the volume enclosed by a convex curved surface will always underestimate the size of the void. As a result, any calculation of volume returned by this method will not be accurate. This is not necessarily a problem as long as the calculation is consistent, but there are other problems that will make this method produce inconsistent measurements as well. The first of these additional problems is that the wound is not flat in a 2-dimensional plane, as the height of the edge varies around the perimeter of the wound. As a result, there will be sections of the wound edge where the height is above or below the plane, which will add inconsistent error to the volume calculation. The second issue that will cause inconsistent measurement is that as the wound heals, the edges of the skin can pull back and make the generated plane lower down in height which would make measurements across days inconsistent, or tissue in the middle of the wound can heap up above the plane formed using the perimeter of the wound. In either case, tissue is above the plane, and that would suggest a negative volume, which is impossible. As a result, the tissue growth volume calculations produced using this method are not consistent or meaningful enough to be used for validation.

Therefore, we developed the second method that interpolates a curved surface that would represent what the surface of the skin would be if there were no wound. This “healthy” surface is then used to calculate the volume of the wound void and determine tissue growth. Using the wound edge pairs found in the edge detection task, a buffer of healthy skin is selected from outside the edges of the wound. Sweeping across the X direction and Y direction individually, a 2-term exponential curve is fit across the wound at each index using the healthy skin buffer as input data. This ultimately generates two sets of curves that represent a predicted surface, one where the curves are drawn vertically across the wound and another where the curves are drawn horizontally. Each of these surfaces is smoothed using a 1 dimensional moving mean filter in the direction opposite the direction that the curves were drawn, then convolved with a 2-dimensional gaussian filter to further smooth out the surface and reduce noise. The two surfaces are then averaged together to produce a final predicted surface that will be used for the volume calculation. The final volume is calculated in the same way as the flat plane enclosed volume; at each point (X,Y) within the wound, the difference in height (Z) between the predicted surface and measured surface is summed together and equals the total volume of the wound void.

## Results

### Data Extraction

Results of alignment based on the femur can be seen in Figure 1.

**Figure 1.**
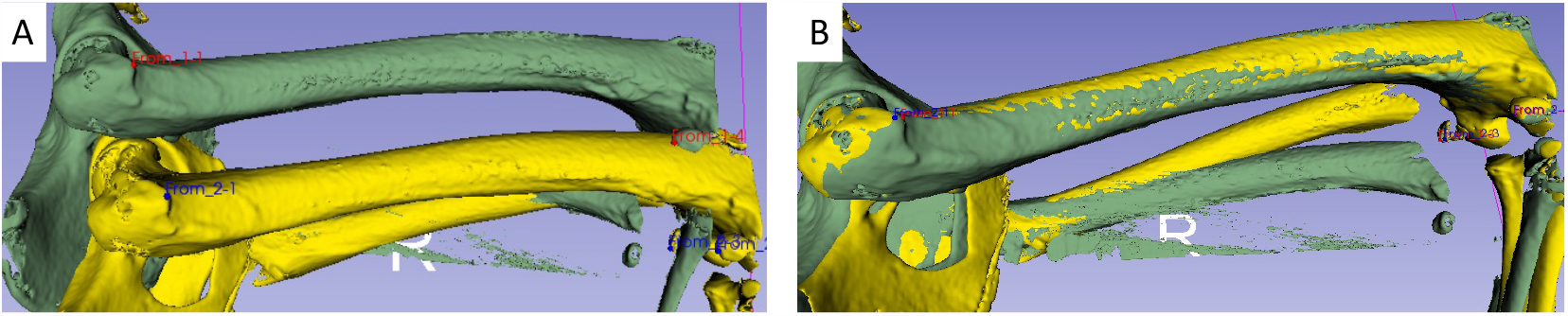
Panel A demonstrates the two scans in their native orientation (Scan 1 = Green, Scan 2 = Yellow). Manual orientation was used to align the two femurs using the Image Guided Therapy (IGT) Package in 3D Slicer. Panel B shows the results of the alignment, Scan 2 (Yellow) superimposed over Scan 3 (Green); green areas visible represent mis-alignment.

The result of the alignment was fairly satisfactory with respect to the femur, with minimal areas of the femur un-aligned. However, given the differential flexion of the knee, due to differences in the position of the dog between the two scans, the lower leg bones are clearly offset. This resulted in a slight mis-alignment with the soft tissue component of the thigh was evaluated (Figure 2). However, since the primary reason of the alignment was to establish a coordinate system for calculating the volume of the wounds, the small amount of misalignment was not considered significant, and would be addressed in the calculation of the wound voids.

**Figure 2.**
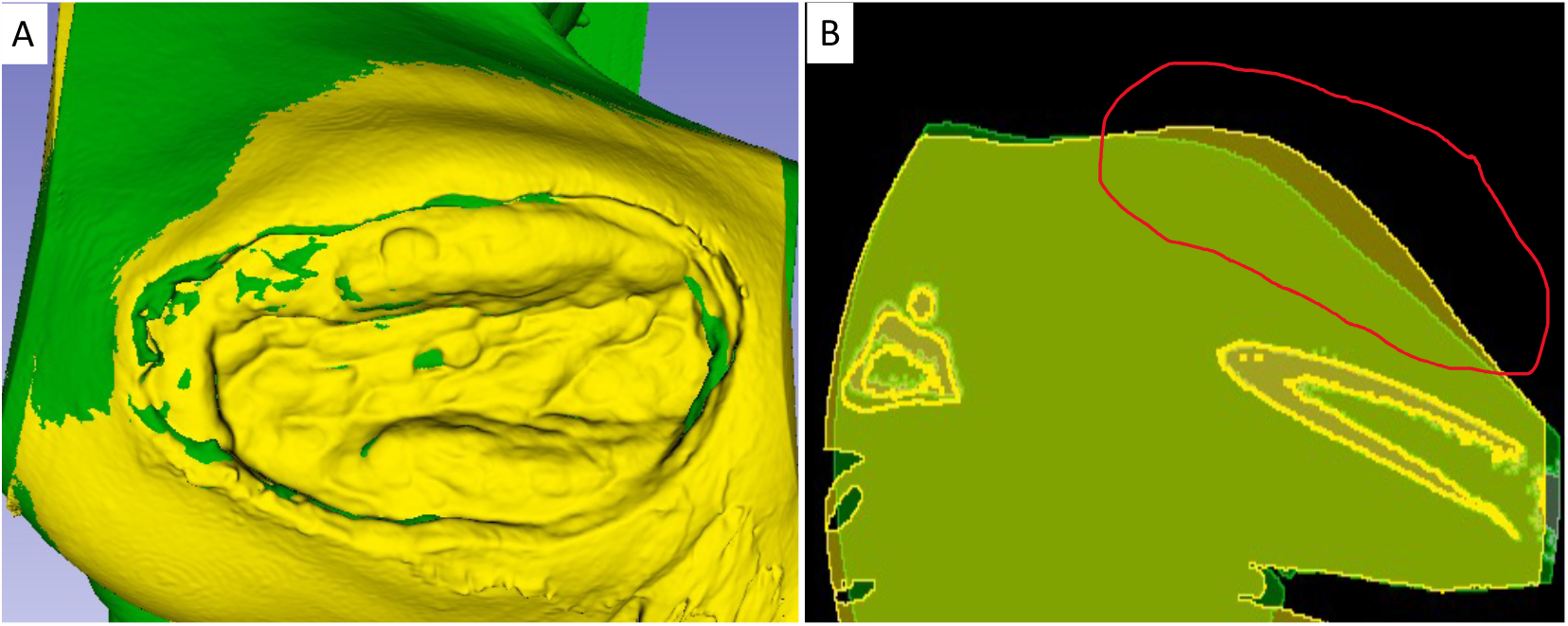
Panel A shows the superimposed soft tissue of Scan 2 (Yellow) over the soft tissue of Scan 1 (Green) based on the alignment process shown in Figure 1. While this alignment appears fairly good, Panel B demonstrates that the soft tissue surface difference even if the femur is well matched (red circled area) Note that this is not actually part of the wound, but rather the adjacent intact tissue. However, this difference will be resolved in the calculation of the wound volumes in the following sections.

### Data Analysis

#### Task 1: Wound Edge Detection

Figure 3 demonstrates the results of wound edge detection. This is a multi-step process, as related in the Methods, and is predicated upon finding a sharp difference in the wound height as measured after alignment to a fixed coordinate system.

**Figure 3.**
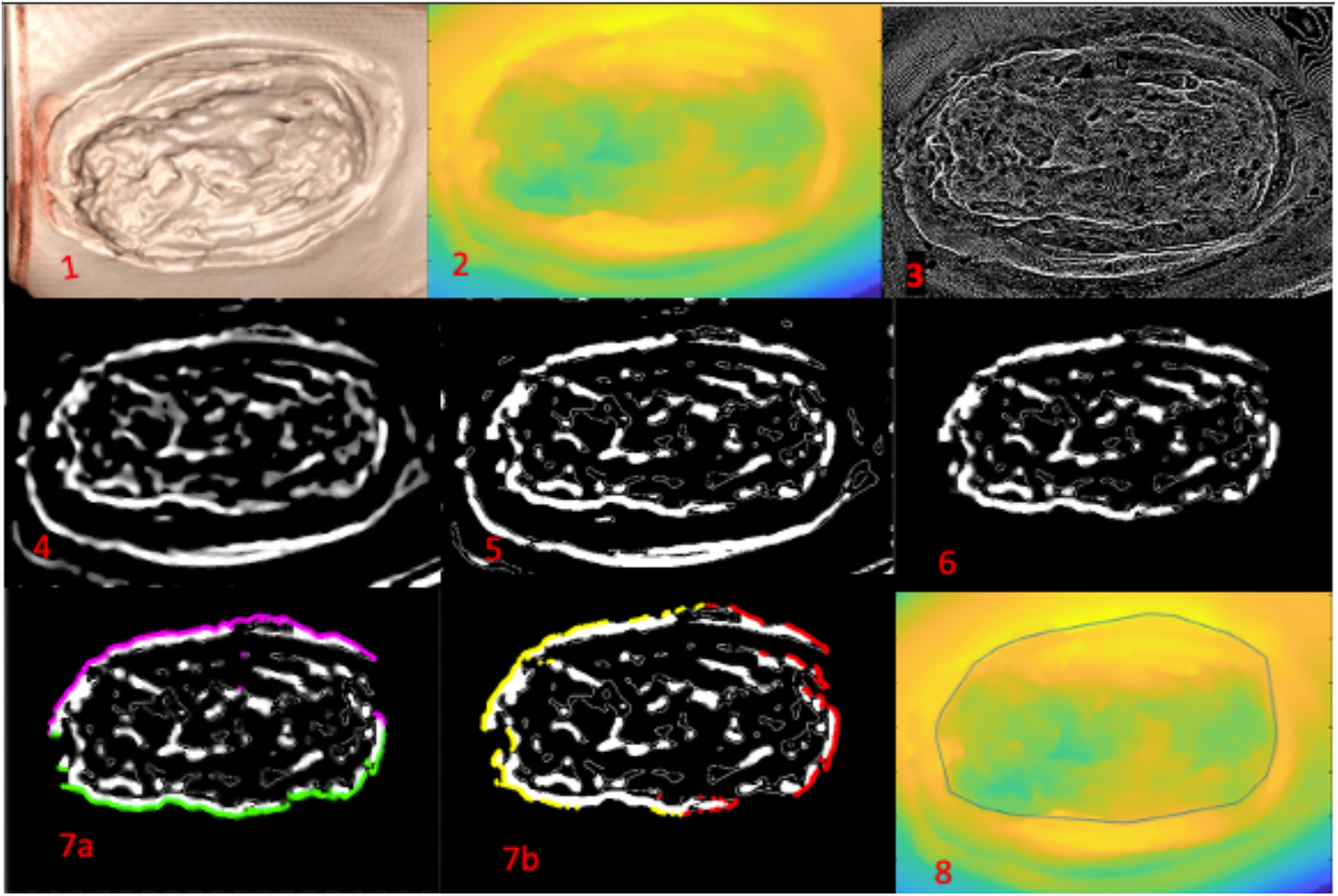
Multiple panels showing sequence of edge detection process. Panel 1 shows the 3-dimensional rendering of the wound generated using 3D Slicer. Pane1 shows a surface height/depth heatmap, where Yellow is the most superficial and blue is the deepest portion of the wound. Note the blue scaling at the lower corners of the panel do not represent the wound, but rather the surface of the dog’s leg as it curves away from the wound. Panel 3 shows a grey-scale depiction of the divergence of surface heightmap. Panel 4 shows the image after a Gaussian filter has been applied and low values rounded to 0. Panel 5 shows the “Sharpen” filter applied and image converted to binary bitmap. Panel 6 shows the wound having been outlined manually and data outside of wound masked. Panels 7a and 7b show marking of the vertical and horizontal edge pairs marked; Panel 7a showing location of top and bottom edge pair, and Panel 7b showing locations of Right and Left edge pair. Panel 8 shows the outline of the wound edge drawn on the heightmap.

#### Task 2 – Wound Volume Determination and Tissue Growth Rate Calculation

Figure 4 depicts the results of Method 1: drawing a flat plane across the edges of the wound and using that as the superficial boundary of the wound void to calculate its volume. The issues with this approach are readily evident. Firstly, the edges of the wound do not exist in a single plane; this results in the need for manual selection of which points of the wound edge are used to generate the flat plane, which introduces inconsistency from scan to scan. The second issue is that the dog’s leg is a curved object, not only would a flat surface approximation will not give accurate results, but it leads to artifacts where, as the wound heals, areas of the base are projected above the superficial plane, resulting in negative void volume, which is not possible. Therefore, we progressed to Method 2: interpolation of the curved surface of the dog’s leg.

**Figure 4.**
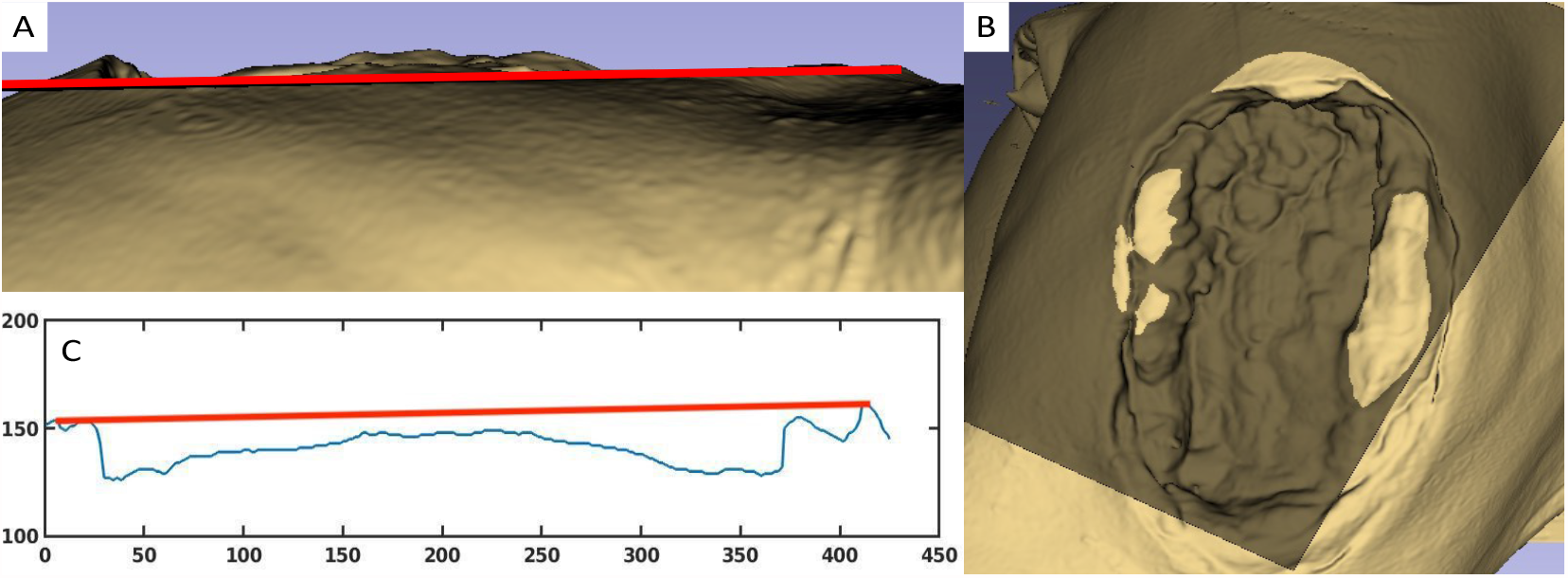
Demonstration of Method 1, drawing a flat plane, to bound the upper surface of the wound. Panel A shows a side view of the wound on Post Injury Day #3 in relation to the projected plane of the upper surface of the wound generated on Day #0 (red line). After alignment to the femur, there are clearly areas of the wound that project above the projected plane, resulting in impossible negative void space. Panel B is a top-down view of the wounds: the tan rendering is the wound on Day #0, with a superimposed wound from Day $3 shown in dark brown. Dark brown shows areas where the Day #3 wound is higher, whereas the tan areas show portions where the Day #0 tissue is higher. Panel C is a graphical representation of the initial plane drawn on Day #0 and shows how the curvature of the leg prevents a straight line from approximating a “healthy” surface accurately. The base of the wound goes almost all the way up to the projected skin plane.

Figure 5 shows the steps of interpolating a curved surface of where a “healthy” skin surface would have been had there not been creation of a wound.

**Figure 5.**
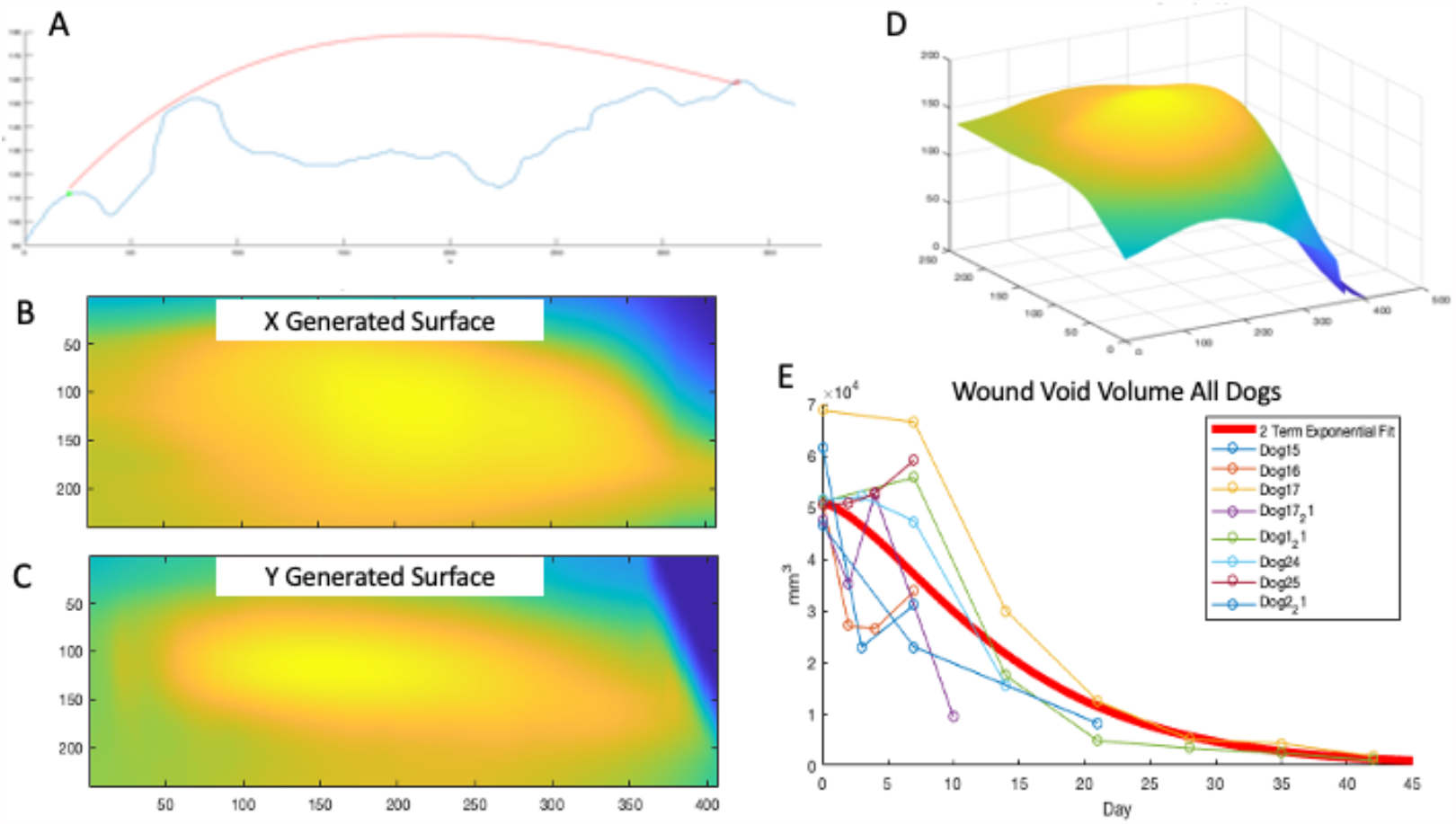
Steps in interpolating curved surface of wound. Panel A shows an example of using the healthy skin up to the edges of the wound to generate an arc to approximate the pre-wound healthy surface (done using Matlab’s curve fitting toolbox). Panels B and C show two raw generated arcs sweeping across the length of the wound (X-axis, Panel B) and across the width of the wound (Y-axis, Panel C); heatmap scale: Yellow = Superficial, Blue = Deep in fixed coordinate system. Panel D shows the final combined surface, smoothed out using convolutional and moving mean filters. Panel E is a graph that depicts plots of void volumes of all wounds for dogs over time with 2 term exponential fit line.

Despite generating a surface that plausibly projects where the healthy skin surface would have been, there are limitations to this method. The first and main limitation is that this method will only produce an approximation of the void volume. By the very nature of generating a predicted surface, it is impossible to get precise measurements that will accurately represent the void. Additionally, this method will not always produce a perfectly smooth and convex surface. This method relies heavily on the skin edges accurately representing the curvature of a healthy leg. In some scans, the bounds of the scan do not fully encapsulate the edges of the wound, and there is not a healthy skin edge to predict from. In these cases, only the surface with curves fit in a single direction can be used to predict the volume, as the other direction will have large error due to poor quality input data. In other scans, as a wound heals and the circumference of the wound contracts, tissue will heap up and change the slope of the skin at the edges of the wound. This natural process will impact the accuracy of the predicted surface and will introduce error in the calculation of the void volume.

However, despite the limitations, the surface interpolation method was found to produce the best results for tissue growth rate measurement with the data available. Due to the deformable nature of soft tissues and the ability for tissue to swell or shrink depending on hydration and inflammation levels, it is impossible to quantify the volume of a tissue void with high precision and accuracy. However, the surface interpolation method is able to produce consistent results of void measurement that can give a quantifiable measure of tissue growth rate, which can then be used to compare different putative interventions and be computed for use as potential reward functions for machine learning/artificial intelligence training.

## Discussion

Volumetric soft tissue and muscle injuries have, by definition, volume, and therefore cannot be effectively characterized merely by measuring surface dimensions. While the use of 3dCT to quantify the volume of various tissue structures is common, these structures invariably are encompassed by other tissue or have clear borders. This is not the case for volumetric soft-tissue or muscle wounds that involve the loss of the skin surface. In these cases, the volume being quantified is a void, which cannot be distinguished from the atmosphere surrounding the individual being scanned. Therefore, the “surface” boundary of the volume of interest cannot be readily identified and needs to be extracted from surrounding tissue features. The deformability of the soft tissue and irregular nature of the wound edges makes a pure algorithmic method extremely difficult to generalize. We have developed a method that merges human visual perception with the quantifying and edge detection capabilities of a software package, 3D Slicer. Being able to quantify the size of a volumetric muscle wound allows the quantification of the rate of healing, which is an crucial capability in the assessment of putative measures intended to speed healing and/or tissue regeneration. The ability to quantify wound size is even more critical if there are to be attempts to use computational methods, such as artificial intelligence systems, to control the healing process. Such systems, which include deep reinforcement learning, rely on optimizing a reward function, which needs to be a computable metric. We hope that the methods presented in this paper will aid in further development to automate the quantification of these types of wounds.

## Author Contributions

Conceptualization, C.C. and G.A.; methodology, D.L., C.C. and G.A.; generation of experimental data, S.J., K.H. and S.B; writing—original draft preparation, D.L and G.A.; writing—review and editing, D.L, C.C., Y.V. and G.A.; visualization, D.L. All authors have read and agreed to the published version of the manuscript.”

